# Local adaptation in climate tolerance at a small geographic scale contrasts with broad latitudinal patterns

**DOI:** 10.1101/2025.05.27.655235

**Authors:** Greg M. Walter, Avishikta Chakraborty, Fiona E. Cockerell, Vanessa Kellermann, Lesley A. Alton, Craig R. White, Matthew D. Hall, Carla M. Sgrò

## Abstract

While climate adaptation is typically quantified across broad gradients, the potential for adaptation to the same environmental variables at small scales is rarely tested. If local-scale environmental heterogeneity can generate patterns of adaptation similar to broad gradients, then we currently underestimate adaptive capacity to global change. We quantified population variation in climate tolerance traits and their plasticity for five populations of *Drosophila melanogaster* from a 3000-km latitudinal gradient, which we contrasted with eight local populations from an environmentally heterogeneous 600×300km area. Population variation in stress tolerance at the local scale was comparable to that across latitude. Consistent with local adaptation, populations from warmer and drier environments showed greater heat and desiccation tolerance, and populations from more predictable environments showed greater plasticity. Climate adaptation at smaller geographic scales can therefore be comparable to adaptation across broad geographic scales. However, patterns of adaptation often changed across geography, which makes predicting responses to global change challenging.

## Introduction

Ongoing global change is creating increasingly stressful and unpredictable environments, which is fragmenting available habitat for mobile species, and increasing the vulnerability of long-lived species and those with low dispersal (Parmesan & Yohe 2003; Parmesan 2006). It is therefore likely that many organisms will need to adapt *in situ* if they are to persist. Rapid adaptation to global change will be more likely if gene flow readily moves adaptive alleles across an environmentally heterogeneous landscape (Sexton *et al*. 2011; Sgrò *et al*. 2011; Willi *et al*. 2022; Sexton *et al*. 2024). Predicting the vulnerability of species then relies on understanding how much adaptive genetic variation is available and how it is distributed among populations across landscapes (Razgour *et al*. 2019; Urban *et al*. 2024). However, we still know little about the scale at which environmental heterogeneity generates and maintains adaptation, which limits our ability to predict the adaptive capacity of species facing global change (Byers 2005; Whitlock 2015; Alvarado *et al*. 2022; Meek *et al*. 2023).

The geographic scale at which adaptation arises across a landscape is rarely considered when assessing vulnerability to global change (Bennett *et al*. 2019). Studies of climatic adaptation typically test for latitudinal clines in stress tolerance traits that reflect adaptation to broad climatic gradients (Huey *et al*. 2000; Hoffmann *et al*. 2002; Etterson 2004; Trotta *et al*. 2006; Sgrò *et al*. 2010; Lasne *et al*. 2018; Ren *et al*. 2020; Lush *et al*. 2024). However, we critically lack tests of whether these same patterns of adaptation emerge at smaller geographic scales (e.g., tens or hundreds of kilometres), where gene flow is more likely to move adaptive alleles among populations and increase the adaptive capacity of populations (Stotz *et al*. 2021; Alvarado *et al*. 2022). While the idea that environmental heterogeneity can generate adaptation at different geographic scales has been around for some time (e.g., Cowling & Pressey 2001; Pressey *et al*. 2007; Sgrò *et al*. 2011), comparative tests of adaptation across spatial scales remain rare because adaptation is typically examined at a single spatial scale or in response to different environmental factors across scales. Studies have shown subtle associations between temperature and local genetic differentiation despite little population structure across latitude (Rumberger *et al*. 2025), and that local adaptation can disrupt latitudinal patterns (Hice *et al*. 2012). However, we lack direct contrasts of climate adaptation across geographic scales for the same environmental variables, which are necessary to determine the spatial scale at which adaptive genetic diversity is generated.

The interplay between dispersal (i.e., gene flow) and selection across geography influences the extent of adaptive genetic differentiation at all geographical scales (Slatkin 1987; García-Ramos & Kirkpatrick 1997; Garant *et al*. 2007; DeMarche *et al*. 2019; Booker 2024). Across geography there are two key spatial scales based on the relative strength of gene flow versus selection. At a broad (latitudinal) scale, distances are greater than typical migration events and even relatively weak differences in selection can create population differentiation (Wright 1943; DeMarche *et al*. 2019). By contrast, at local scales, defined as distances where substantial gene flow occurs (within hundreds of km for highly dispersive species, such as *Drosophila melanogaster*; Gockel *et al*. 2001), local adaptation can only arise when strong differences in selection overcome the homogenizing effects of gene flow (Slatkin 1987; García-Ramos & Kirkpatrick 1997; Lenormand 2002; Richardson *et al*. 2014). Beyond these differences in the balance between gene flow and selection, the type of selection can also differ across geography.

Adaptation along latitude arises in response to broad abiotic gradients, particularly gradual changes in temperature and seasonality that are correlated and create consistent directional selection across large distances (Endler 1977; Sunday *et al*. 2011). By contrast, environmental heterogeneity at a local scale can be created by microhabitat variation, topographic complexity, or localized climatic conditions that create spatial mosaics of selection within the gene flow neighbourhood (Garant *et al*. 2007; Richardson *et al*. 2014). Changes in directional selection at local scales can then make it difficult to predict patterns of adaptation at local versus broad scales (Lenormand 2002). Moreover, when populations face multiple environmental stressors, adaptation to one stress may constrain adaptation to others (Agrawal 2020; Willi & Van Buskirk 2022). For example, evolving high heat tolerance may limit the capacity to adapt to desiccation stress. Given that climate change imposes multiple simultaneous stressors (Urban *et al*. 2024), understanding geographic patterns of adaptation requires examining responses across a suite of climate-related traits.

In addition to adaptation, plastic changes in climate tolerance traits also help populations cope with global change (Reed *et al*. 2011; Gunderson & Stillman 2015; Magozzi & Calosi 2015; Sgrò *et al*. 2016). Environments that are variable and predictable favour the evolution of greater plasticity (Scheiner 1993; Leung *et al*. 2020), which creates populations that differ in their magnitude and direction of plasticity when they adapt to different environments (Lind & Johansson 2007; Murren *et al*. 2014; Matesanz & Ramírez-Valiente 2019; Walter *et al*. 2022). Lower plasticity but greater overall climate tolerance is then expected to evolve in environments that are either consistently harsh, or variable but unpredictable (Reed *et al*. 2010; Botero *et al*. 2015). Both adaptation and plasticity can therefore vary across spatial scales (and interact with each other) to determine population responses to environmental change. At a local scale, fine-grained environmental heterogeneity can favour plasticity that allows individuals to cope with variability within their lifetime (Baythavong 2011). By contrast, broad environmental gradients may favour greater overall climate tolerance at lower (constantly warm) latitudes but greater plasticity at higher latitudes where temperatures are more variable (Janzen 1967; Ghalambor *et al*. 2006). These scale-dependent patterns in the balance between adaptation and plasticity in climate tolerance remain largely untested but are critical for predicting population responses to global change.

To contrast the extent of climate adaptation to the same environmental variables across geographic scales, we focus on geographic variation in populations of the vinegar fly, *Drosophila melanogaster*. Since its introduction to the east coast of Australia, c.100 years ago, *D. melanogaster* has developed latitudinal clines in numerous traits, including body size and thermal tolerance (reviewed in Hoffmann & Weeks 2007; Lasne *et al*. 2018). Molecular data suggest that population structure is weak along latitude due to some level of gene flow at distances up to 1500km (Gockel *et al*. 2001; Kennington *et al*. 2003). Such broad patterns of adaptation and high potential for gene flow mean that *D. melanogaster* provides a powerful conservative test of whether small-scale geographic variation can generate and maintain local adaptation. This is because local adaptation should only emerge if differences in selection at local scales can outweigh the high connectivity present among populations of *D. melanogaster*. We sampled five populations of *D. melanogaster* across 3000-km in latitude along east coast Australia (18°S–43°S), which we compared to eight populations sampled from a variety of climates within a 600km×300km area in Victoria (**Fig. 1a**). Compared to the latitudinal populations, locations sampled in Victoria showed greater fluctuations in temperature, both across and within days (**Fig. 1b-c**). Using a laboratory experiment, we tested how populations across geography varied in climate tolerance traits, as well as in their plasticity to temperature treatments.

**Fig. 1.**
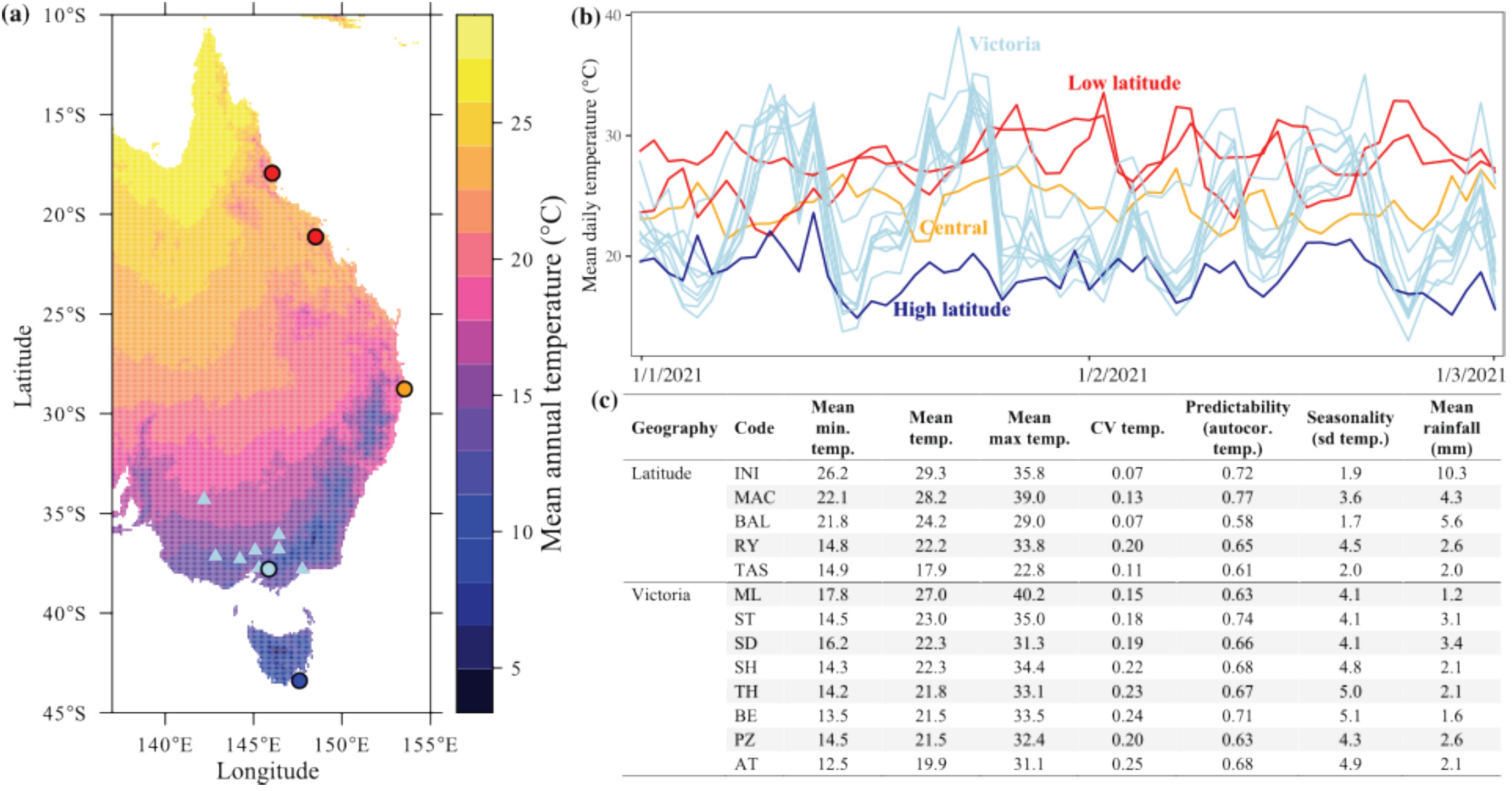
**(a)** Map of sampling locations along east coast Australia (circles; n=5) compared to locations within Victoria (triangles; n=8). Shading represents the average mean annual temperature, where lighter shades (warmer colours) depict warmer areas. **(b)** Mean daily temperature for the sampled locations. Victorian populations show larger fluctuations in temperature. **(c)** Summary of temperature (temp.) and rainfall data for each sampled location across four months prior to sampling. Mean refers to daily mean for each of the variables. Data produced using *NicheMapR* (Kearney & Porter 2020).

We test three hypotheses to contrast climate adaptation across geography: (I) We would detect variation among populations in Victoria for climate tolerance traits and their plasticity, but variation across latitude would be greater. (II) If among-population variation is adaptive, tolerance traits would be associated with the climate of origin. (III) Population variation in thermal plasticity would be associated with environmental variation and predictability. If local environmental heterogeneity generates adaptive differentiation that approaches what is observed across latitude, we may be underestimating the adaptive capacity of local populations facing global change. Adaptation across latitude should generate higher heat tolerance, lower cold and desiccation tolerance, and smaller body sizes at lower latitudes. More constant higher temperatures at low latitudes should generate lower plasticity (Walter *et al*. 2025). If local environmental heterogeneity promotes adaptive differentiation within Victoria, variation among populations should mimic latitudinal patterns: populations from warmer, less variable environments should resemble lower-latitude populations. However, because warmer environments in Victoria are more variable, greater thermal plasticity could instead emerge and contrast with latitudinal trends.

## Methods

### Collection and maintenance of experimental populations

To ensure our estimates of among-population variation would be conservative, we sampled flies in Autumn, which is after the summer peak population size that has the greatest potential for homogenization of populations via dispersal (Shpak *et al*. 2010; Behrman *et al*. 2015). In March–May 2021, we collected 20–50 field-inseminated female *D. melanogaster* from thirteen locations in Australia using sweep nets over fruit waste. For the five latitudinal populations, we sampled flies at fruit farms in Tasmania, New South Wales, Victoria and Northern Queensland (**Fig. 1a**; **Table S1**). In Victoria, we sampled flies from discarded grape skins at eight wineries from a variety of climates (**Fig. 1a**; **Table S1**). We chose wineries due to the abundance of breeding habitat at the same time of year, allowing us to sample broadly across the state within a short period. For each population, we generated a mass-bred population in the lab from 19-60 isofemale lines (mean=44.6; **Table S1**), which we maintained at large sizes (1000-1500 individuals) in five 300mL bottles. We kept all populations at 25°C for 12 generations (c.9 months) to remove transgenerational environmental effects and focus on genetic differences among populations, and to ensure sufficient mixing of genotypes to break up linkage disequilibrium (Kellermann *et al*. 2015). **Methods S1** details population establishment and maintenance.

### Manipulative experiment

In January 2022, we conducted a laboratory experiment by rearing larvae from all 13 populations under three temperature treatments (13, 25 and 29°C), and measuring wing size and tolerance traits (heat, cold and desiccation) on emergent adults. 25°C represents control conditions, and 13 and 29°C represent the cold and hot margins that larvae in natural populations are often exposed to in winter and summer, respectively. For each population, we placed c.200 adult flies in each of two laying pots with a c.3mm layer of food and let them lay for 6 hours. We then transferred 20 eggs into each vial (n=36–42 vials/population/treatment; **Table S2**) containing 7mL of food and left larvae to develop in their treatment. To keep replication feasible, we split the experiment into two temporal blocks so that adults emerged 21–28 January (block 1) and 1–8 March (block 2). For each block, we conducted 2–3 rounds of egg-picks for each treatment over 4–5 days to maximise the overlap in eclosion across treatments. We timed the egg picks according to pilot experiments so that adult flies would emerge simultaneously. We used 1456 vials in total (n=455–546 vials/treatment).

Eclosed adults from each treatment were collected in separate vials each day so that flies would be the same age across treatments. Flies were kept at their experimental treatments until they reached maturity and mated. Pilot experiments suggested flies kept at 13°C would not mature, and so they were kept at 16°C to mature in time for the experiment.

### Stress tolerance assays and body size measurements

We measured females, which we collected under light CO_2_ anaesthesia, let them recover (48 hours) and then used established methods to measure heat, cold, or desiccation tolerance, or body size (Hoffmann *et al*. 2002; Lasne *et al*. 2018). We measured 30 females per population in each assay, which we split across the two blocks. Different individuals were measured in each assay. To assay heat tolerance, we measured heat knockdown by putting individual females into sealed 5mL glass vials, submerging them in a 39°C water bath and measuring the time for each female to cease moving. To assay cold tolerance, we measured chill-coma recovery by placing individual females into sealed 1.5mL plastic tubes that we submerged in a 0°C water bath for 4 hours and measured the time they took to regain movement. To measure desiccation tolerance, we put individual females in unsealed 5mL vials placed in sealed glass tanks containing silica gel (to maintain relative humidity <5%) and recorded the time they took to cease moving. To measure wing size, we removed the right wing of 30 females, mounted them on a microscope slide and photographed them (M60 stereo microscope, Leica, Heerbrugg, Switzerland). We used *imageJ* to quantify 10 vein intersection landmarks (**Fig. S1**) and estimate centroid size.

### Statistical analyses

We used *R* (v.4.3.2) statistical software for all analyses (R Core Team 2024). We applied linear mixed models using *glmmTMB* (Brooks *et al*. 2017) and checked model assumptions using *DHARMa* (Hartig 2022). We tested for significant interactions using type-III ANOVA.

#### Hypothesis I

##### Testing for among-population variation in tolerance traits and body size

To test whether variation among populations was significantly different across geography, we applied the linear mixed model

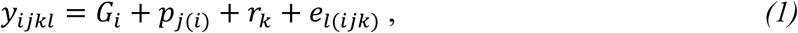

where geography (*G_i_*; latitude vs Victoria) is a fixed effect. Replicate run nested within block was a random effect (*r_k_*) and *e*_*l*(*ijk*)_ is the residual. We applied equation 1 to each trait and treatment temperature separately. We included population (*p*_*j*(*i*)_) as a random effect and used log-likelihood ratio to test the null model that estimates a single variance among populations and so assumes equal variance across regions, i.e., *p*_*j*(*i*)_∼*N*(0, *σ*^2^), with the alternative model estimating an among-population variance for each geographical region, and so 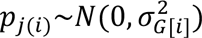 where *i* indexes region. Significant differences between models suggest that among-population variance differs between latitude vs Victoria.

Next, we tested whether variation among populations was significant within each region, and whether populations (within region) varied significantly in their response to temperature treatments. We used equation 1, but analysed latitudinal and Victorian populations separately, included population and temperature as fixed effects and tested for significant population×temperature interactions. Replicate run was the only random effect.

#### Hypotheses II

##### Testing whether among-population variation is adaptive

For all 13 locations, we extracted fine-scale rainfall (daily) and temperature (hourly) data using *Nichemapr* (Kearney & Porter 2020) for the three summer months preceding sampling. To quantify temperature variation, we calculated standard deviation across the three months. To quantify temperature predictability, we used the *acf* function to calculate autocorrelation of daily mean temperature across the three months. To test for associations between climate of origin and assayed traits, we used

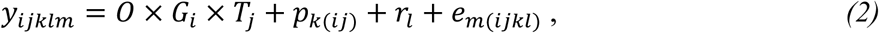

where the three-way interaction of the fixed effects (*O* × *G_i_* × *T_j_*) includes environment of origin (O) measured at the population locations (continuous variable), the geographical comparison (*G_i_*; Victoria vs latitude) and treatment temperature (*T_j_*). Random effects included the *k*th population nested within treatment temperature and geography (*p*_*k*(*ij*)_) to avoid pseudoreplication, and replicate block (*r_l_*). *e*_*m*(*ijkl*)_ is the residual. We applied equation 2 to each trait separately. We included mean temperature as the environmental covariate (O) for analysing thermal tolerance and body size, and mean daily rainfall for desiccation tolerance.

A significant slope for O (effect of environment) provides evidence that environment of origin is associated with that particular trait. A significant two-way environment of origin×geography interaction suggests that latitudinal and Victorian populations differ in their association between the environment of origin and trait. A three-way interaction then suggests the association between environment and trait changes with geography and treatment.

#### Hypothesis III

##### Testing whether environment determines plasticity

To compare differences in plasticity, we extracted the population marginal means from the second application of equation 1 using *emmeans* (Lenth 2019), and then calculated plasticity using: 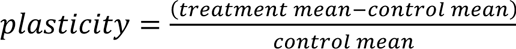, which represents plasticity as the proportional change in phenotype from the 25°C control treatment. This removes population differences in the trait to focus on estimating plasticity of each population (Valladares *et al*. 2006). We then used equation 2 with environmental variation and predictability traits as the covariate, and plasticity as the response variable.

## Results

### Hypothesis I

#### High variation among Victorian populations for tolerance traits and plasticity

Only body size showed greater variation across latitude compared to among Victorian populations (**Fig. 2**; **Table 1a**). For all tolerance traits, variation among populations was not significantly different for latitudinal versus Victorian populations (**Fig. 2**; **Table 1a**). The magnitude of differences among Victorian populations was therefore similar to that observed across latitude for heat and cold tolerance, and consistently larger (but non-significant) than latitudinal populations for desiccation resistance. Latitudinal populations showed >3-fold greater differences than Victorian populations for wing size (**Table 1b**).

**Fig. 2.**
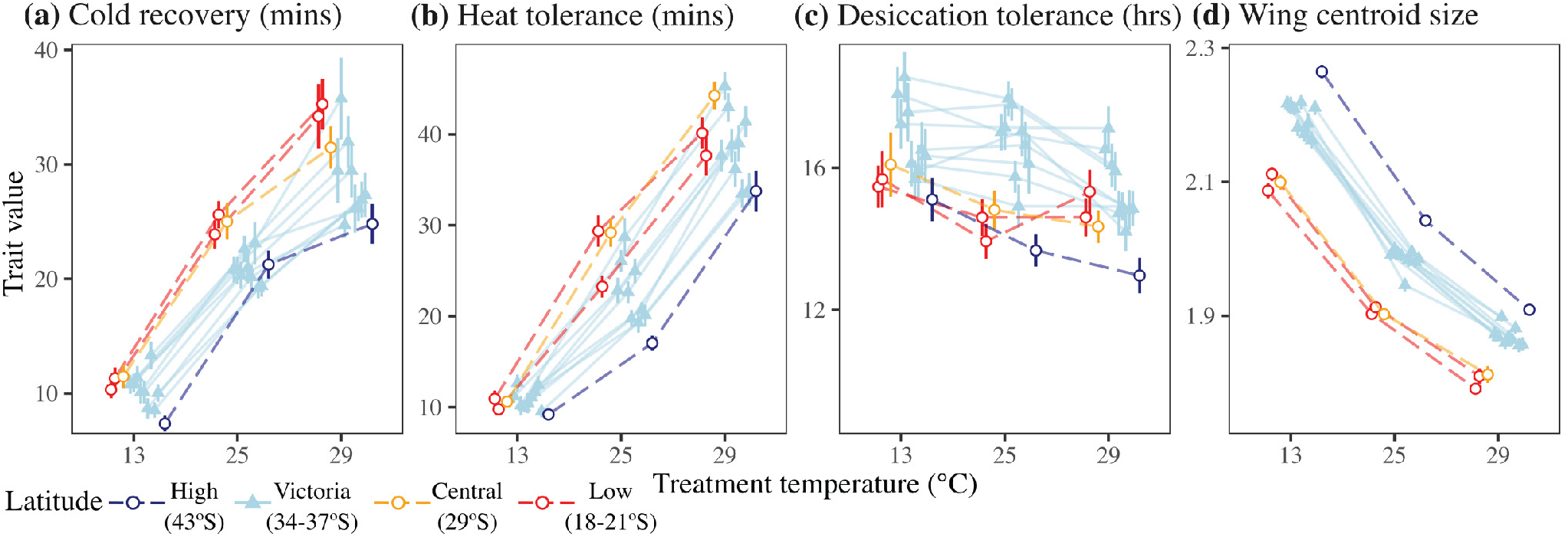
The effect of treatment temperature on stress tolerance and body size for populations along latitude (open circles) and within Victoria (closed circles). We measured **(a)** cold tolerance as chill coma recovery time (mins), **(b)** heat tolerance as heat knockdown time (mins), **(c)** desiccation tolerance (hours), and **(d)** body size as wing centroid size. Points and error bars represent the marginal means (±1SE). Data points are offset for clarity.

**Table 1.**
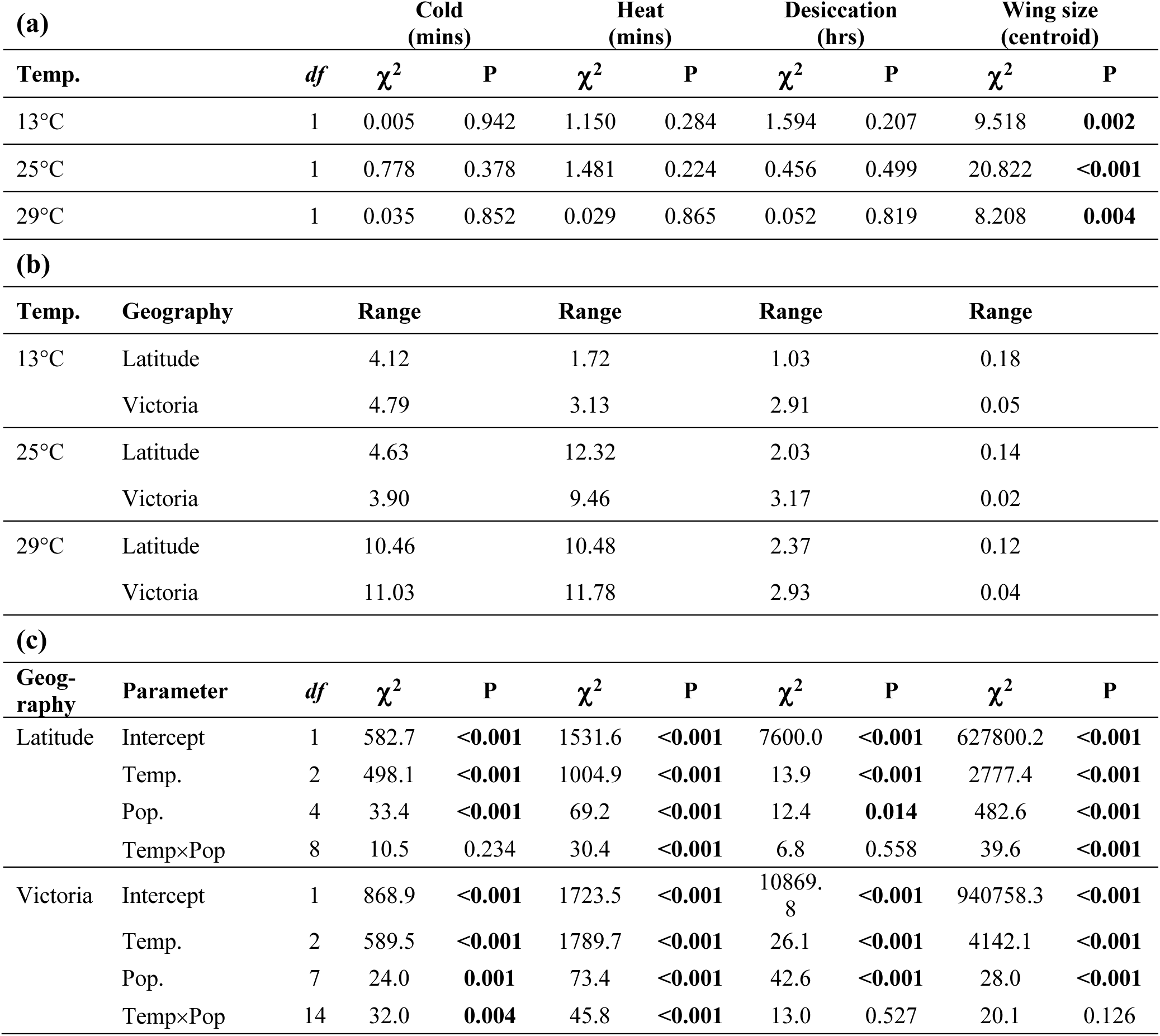
Testing for differences in among-population variation across geography both within and across temperature treatments. **(a)** Likelihood-ratio tests (equation 1) assessing differences in variation among populations for latitudinal versus Victorian populations (within each treatment temperature). **(b)** Effect sizes for maximum differences among populations for latitudinal and Victorian populations, which were calculated as the difference in marginal mean between populations with the largest and smallest values for each trait (i.e., the range). **(c)** ANOVA summaries testing population×temperature interactions within each geographic region. Parameters in bold are significant at p<0.05. Temp = treatment temperature, and Pop = population.

Temperature created significant differences in each trait. Flies exposed to colder temperatures were larger, more cold tolerant and less heat tolerant (**Fig. 2**). Within each geographic scale, we found significant variation among populations in all tolerance traits and body size (**Table 1c**). Statistically significant interactions between population and temperature quantifies how populations vary in response to treatments, and therefore reflects population variation in plasticity. Both Victorian and latitudinal populations displayed significant variation in plasticity, but for different traits (**Fig. 3**). Victorian populations only showed significant temperature×population interactions for cold and heat tolerance, whereas latitudinal populations showed significant interactions for heat tolerance and wing size (**Table 1c**). Desiccation tolerance showed no significant interaction.

**Fig. 3.**
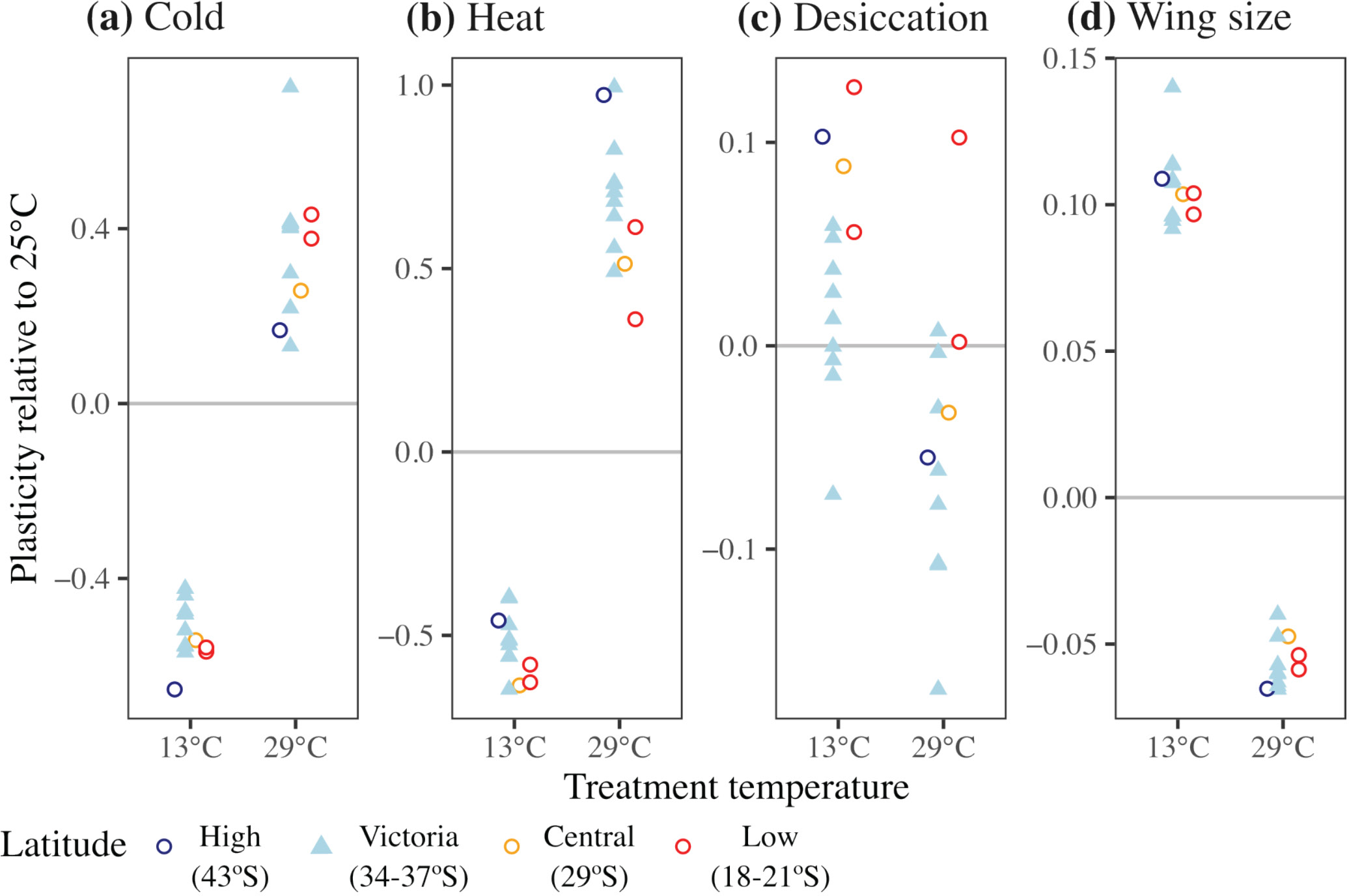
Population variation in thermal plasticity for each trait (**a-d**). Triangles represent Victorian populations, and circles the latitudinal populations. Horizontal grey lines represent the control conditions (25°C), and the points represent the plasticity for each population as the change in phenotype from the control condition. Negative and positive values then represent a trait decrease and increase, respectively, relative to the 25°C treatment. Data points are offset for clarity.

##### Differences in climate across geography

While temperatures were higher and more consistent at lower latitudes, Victorian locations experienced greater variation in temperature across days, and within each day (**Fig. 1c**). Rainfall was lower at high latitudes, ranging 2–10mm/day south–north, while Victorian locations varied between 1.2-3.4mm/day (**Fig 1c**). Correlations between mean temperature and both rainfall and temperature predictability were stronger across latitude (0.81, 0.84) than within Victoria (−0.06, −0.37) (**Fig. S3**).

### Hypothesis II

#### Population variation was associated with climate of origin

To test whether among-population variation in tolerance traits and body size was adaptive, we quantified associations between climate of origin with the traits measured in the laboratory. Below we summarise the results for each trait.

##### Cold tolerance

We found a significant three-way interaction between temperature of origin (O), geography (G; latitude vs Victoria) and treatment temperature (T; **Fig. 4a**; **Table S3**). While there was no association between temperature of origin and cold tolerance at 13°C, at 25°C and 29°C, latitudinal populations from warmer locations showed slower cold recovery (reduced cold tolerance), as expected (**Fig. 4a**). By contrast, Victorian populations showed the opposite trend where populations from warmer environments showed higher cold tolerance (**Fig. 4a**), suggesting that they were better at coping with a cold shock after acclimating to warmer conditions.

**Fig. 4.**
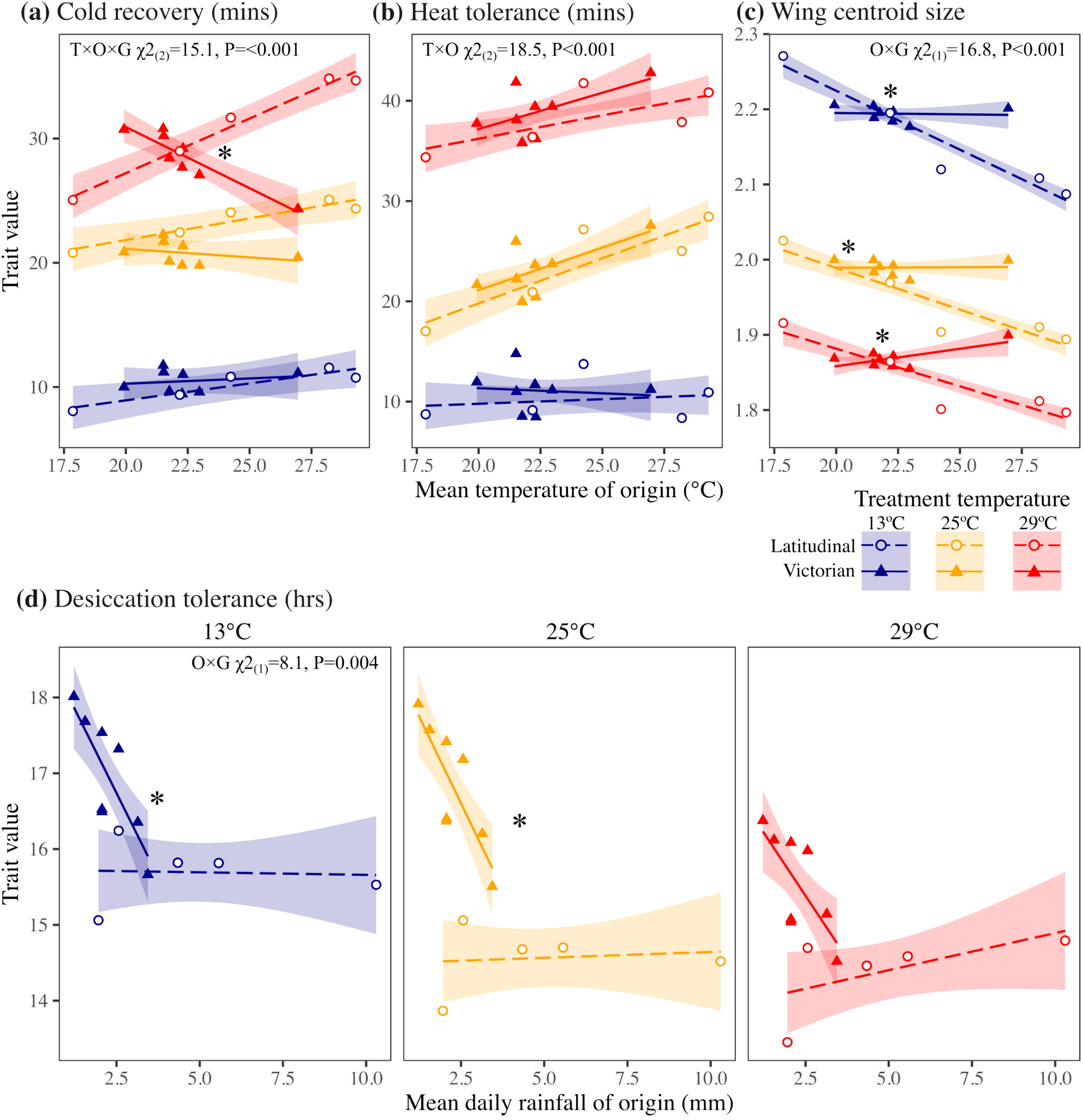
Association between climate of origin and each trait (**a–d**) after exposure to three temperatures (different colours). Panels **a–c** compare mean daily temperature to cold tolerance, heat tolerance and wing size, while panel **d** compares average daily rainfall to desiccation tolerance. In panel **d**, treatment temperatures are presented in different panels to aid interpretation as they otherwise overlap substantially. Shading represents ±1SE around the regression estimates. Significant interactions are presented at the top of each panel, where T=treatment temperature, O=environment of origin and G=geographical comparison (latitude vs Victoria) – **Table S3** contains the full ANOVA summaries. Asterisks represent significant differences in slopes between Victorian and latitudinal populations.

##### Heat tolerance

We found no significant effect of geography, suggesting that associations between temperature and heat tolerance were consistent for Victorian and latitudinal populations (**Fig. 4b; Table S3**). While there was no strong association between temperature of origin and heat tolerance at 13°C, populations from warmer locations showed higher heat tolerance in warmer treatments (**Fig. 4b**), creating a positive association supported by a significant temperature of origin×treatment interaction (**Table S3**). We therefore found evidence of local adaptation in warmer treatments, which was consistent for Victorian and latitudinal populations.

##### Wing size

A significant temperature of origin×geography interaction suggests that wing size showed contrasting associations with temperature of origin for Victorian versus latitudinal populations (**Fig. 4c; Table S3**). While populations from warmer lower latitudes had smaller wings (consistent with latitudinal studies), Victorian populations from warmer locations either had slightly larger wings (29°C treatment) or no association between temperature of origin and wing size (13°C and 25°C; **Fig. 4c**).

##### Desiccation tolerance

Latitudinal populations showed weak associations between rainfall and desiccation tolerance, whereas Victorian populations from drier environments showed greater desiccation tolerance that was consistent across treatment temperatures (**Fig. 4d**). This was supported by a significant rainfall of origin×geography interaction (**Table S3**). Victorian populations from drier environments therefore showed greater desiccation tolerance, suggesting local adaptation to water availability.

To ensure that body size did not affect the association between climate of origin and each trait, we conducted the same analyses, but used population means and included wing size as a covariate. This did not change the strength of the association between climate of origin and tolerance traits, suggesting the patterns observed were not due mainly to variation in body size.

### Hypothesis III

#### Heat tolerance plasticity is linked to environmental predictability

To test for an association between environmental variation and plasticity, we focussed on heat tolerance as it showed larger variation in plasticity among populations compared to other traits (**Fig. 3**) and the most consistent association with climate (**Fig. 4**). In the hot treatment, greater predictability of temperature across days was associated with larger increases in heat tolerance for the Victorian populations, but smaller increases in heat tolerance along latitude (**Fig. 5a**). This was supported by a significant interaction between autocorrelation of mean temperature and geography (**Table S4**). Although non-significant in the cold treatment where overall heat tolerance was lower, greater predictability was associated with smaller decreases in heat tolerance in Victorian populations, but no relationship for latitudinal populations (**Fig. 5a**). We found no association between the amount of temperature variation and plasticity (**Fig. 5b**). Victorian populations from more predictable thermal environments therefore showed more adaptive plasticity – greater increases in heat tolerance in the hot treatment, smaller reductions in heat tolerance in the cold treatment (**Fig. 5**). By contrast, latitudinal populations showed weaker, opposing trends.

**Fig. 5.**
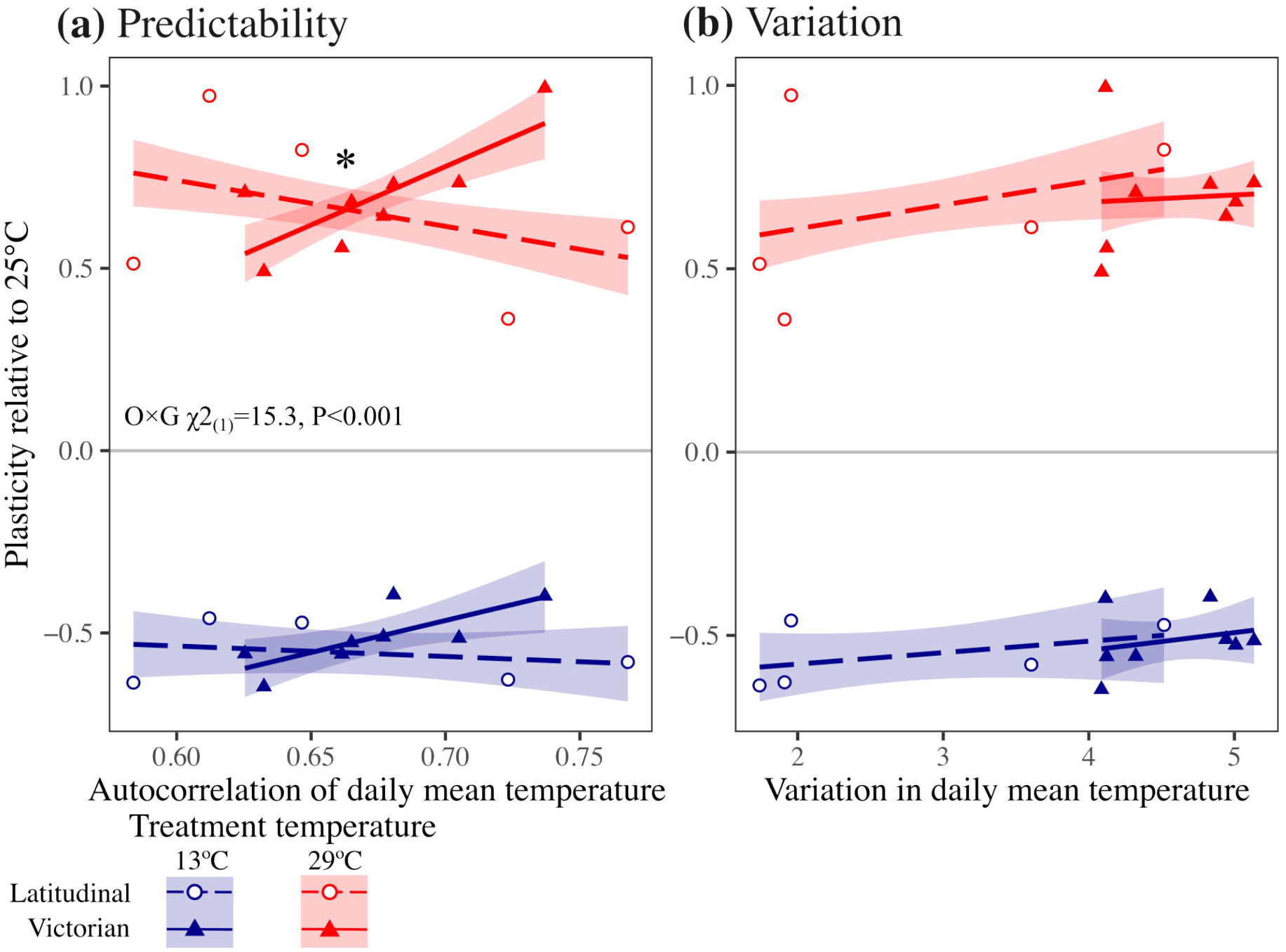
Association between **(a)** predictability and **(b)** variation of daily mean temperature across three months, with plasticity in heat tolerance. Temperature predictability is the autocorrelation of mean daily temperature across the three months preceding sampling, while temperature variation was calculated as the standard deviation of temperature across the three months. Shading represents ±1SE around the regression estimates. Triangles represent Victorian populations, and open circles the latitudinal populations. The significant O×G interaction is presented, where O=environment of origin and G=geographical comparison (latitude vs Victoria). This provides evidence that Victorian and latitudinal populations differ in their relationship between autocorrelation in temperature and thermal plasticity – **Table S4** contains the full ANOVA summaries. Asterisks represent significant differences in slopes between Victorian and latitudinal populations.

## Discussion

Using populations of *Drosophila melanogaster*, we contrasted population variation in tolerance traits and body size across latitude with population variation at a smaller geographical scale within Victoria. Among-population variation in Victoria was significant for all traits (except wing size), and similar in extent to that seen across latitude (*Hypothesis I*: **Figs. 2–3**). As evidence for local adaptation across Victoria, population variation in heat and desiccation tolerance was associated with the climate of origin (*Hypothesis II*: **Fig. 4**). Population variation in heat tolerance plasticity was more strongly associated with temperature predictability across Victoria than across latitude, which showed weaker opposing trends (*Hypothesis III*: **Fig. 5a**). Together, our results suggest that environmental heterogeneity to the same environmental variables at a relatively small geographic scale can generate local adaptation to climate that is comparable to the extent of latitudinal adaptation. Local-scale climatic heterogeneity can therefore generate adaptive differentiation that can increase the adaptive capacity of the metapopulation. However, latitudinal and Victorian populations often showed contrasting patterns of adaptation (**Figs. 4–5**), suggesting that predicting population responses to global change needs to consider how adaptation arises across geography.

### Contrasting patterns of adaptation across geographic scales

We show that adaptive divergence among populations can occur at a smaller scale than previously anticipated, which increases the adaptive capacity of the metapopulation to global change. Given that gene flow increases at smaller spatial scales (Sexton *et al*. 2024), populations could rely on locally segregating adaptive alleles rather than their movement across broad geographic scales (Meek *et al*. 2023; Sexton *et al*. 2024). However, our results raise several concerns about predicting population responses to global change based on patterns of adaptation across geography. First, associations between climate of origin and all traits (except heat tolerance), as well as plasticity in heat tolerance, differed for Victoria populations compared to latitudinal populations. Inferences for natural populations facing global change are therefore likely to change with geography. Second, associations between climate of origin and thermal (hot and cold) tolerance changed with treatment temperature, suggesting patterns of adaptation were not only specific to geography, but also the treatment in which they were measured.

### Adaptive differentiation across a heterogeneous landscape

Few studies have directly contrasted climatic adaptation across geographic scales to the same environmental variables. In lodgepole pines, regional variation accounted for 20% of total genetic variance in key ecological traits, suggesting that environmental heterogeneity maintains genetic variation (Yeaman & Jarvis 2006). At a molecular level, subtle adaptive genetic differences in a sea urchin at a local scale (<100km) were associated with temperature despite little population structure across latitude (Rumberger *et al*. 2025). While Hoffmann *et al*. (2001) found evidence of variation among populations for cold tolerance in *D. melanogaster* at local scales (c.20-100km), they did not test heat tolerance or connect among population variation to the environment of origin. Nadeau and Urban (2024) demonstrated variation in thermal tolerance among *Daphnia* from nearby pools, but this variation was not associated with environmental differences, suggesting that microgeographic variation did not generate patterns of adaptation. While these studies suggest among-population variation in climate adaptation can be generated at local scales, they either struggled to connect such variation with environmental heterogeneity, or link adaptive genetic variation to key climate-tolerance traits. By contrasting populations across latitude with local populations from variety of climates, we showed how environmental heterogeneity in the same environmental variables generates adaptive trait divergence in climate tolerance at different geographical scales.

### Predictions about adaptation change across geography

Our key result is that understanding climatic adaptation changes with environments across geography. While Victorian populations mimicked latitudinal heat tolerance trends (populations from warmer areas showed higher heat tolerance), this was not the case for any other tolerance trait or body size. Populations from lower (warmer) latitudes showed higher heat tolerance, lower cold tolerance and smaller body size (**Fig. 4a-c**), suggesting that thermal tolerance and body size are selected together across latitude due to strong correlation of environmental variables (**Fig. S3**) and/or latitudinal trends in genetic factors (e.g., inversions; Rako *et al*. 2006). By contrast, different combinations of traits could be favoured in Victoria, away from the coast. While Victorian populations from warmer locations showed higher heat tolerance (matching the latitudinal pattern), they differed from latitudinal trends by showing higher cold tolerance and larger body sizes (**Fig. 4**), and stronger associations between moisture availability and desiccation resistance (**Fig. 4d**). Although latitudinal clines are a powerful tool for understanding adaptation to broad climatic gradients, they likely represent adaptation to a specific combination of environmental variables. Broad patterns of adaptation are then likely to provide inaccurate predictions for population responses to global change when applied to different geographic scales or environments (O’Brien *et al*. 2017; Cornwell *et al*. 2021; Sexton *et al*. 2024; Urban *et al*. 2024).

### Temperature-dependent patterns of adaptation

Compared to latitudinal populations, patterns of adaptation in thermal tolerance changed across treatment temperatures to a greater extent for Victorian populations (**Fig. 4**). The cold treatment tended to reduce variation among populations in thermal tolerance, whereas the hot temperature increased variation among populations. However, greater variation at the high temperature weakened patterns of adaptation in heat tolerance (**Fig. 4b**), and reversed the expected pattern for cold tolerance – populations from warmer environments were more cold-tolerant (**Fig. 4a**). Across a species range, some populations could therefore be more sensitive to temperature changes than others (Walter *et al*. 2020). Our finding that patterns of adaptation are weakened or reversed under stressful temperatures is consistent with other studies (Booker 2024; Walter *et al*. 2025), potentially increasing vulnerability to climate change if, for example, warm-adapted populations are as vulnerable to novel high temperatures as all other populations (Anderson & Wadgymar 2020). Accurately predicting the response of natural populations to global change therefore requires considering how locally adapted populations will vary in their response to the particular conditions created by environmental change.

### Adaptive differentiation in plasticity across Victoria but not across latitude

While variation or predictability in temperature was weakly associated with plasticity for latitudinal populations, Victorian populations from locations where temperature was more predictable across days showed greater thermal plasticity in heat tolerance, but only in the hot treatment (**Fig. 5**). Given that Victorian populations experience greater variation in temperature than latitudinal populations, these results support theory and empirical observations that variable and predictable environments favour adaptive plasticity (Reed *et al*. 2010; Diaz *et al*. 2020; Leung *et al*. 2020; Cicchino *et al*. 2024; Gallegos *et al*. 2024), which helps populations adapt to thermal variation. Local adaptation across a heterogeneous landscape can therefore generate variation in plasticity, which could determine population responses to environmental change (Hamann *et al*. 2016; Vinton *et al*. 2022). While such adaptive plasticity can help populations buffer familiar thermal variation, it could also increase their vulnerability to unpredictable changes in temperature created by global change (Ashander *et al*. 2016). Understanding how adaptive plasticity evolves across a heterogeneous landscape, and then determining the consequences for persisting under environmental change remains challenging for predicting population responses to global change (Meek *et al*. 2023).

### Limitations

While we show that it is possible for adaptive differentiation to occur at smaller geographic scales than originally anticipated, it is important to interpret our results cautiously. Further work is needed to test whether local adaptation can emerge within regions at different latitudes, or whether it is unique to the greater environmental heterogeneity present in Victoria. Hoffmann *et al*. (2001) quantified cold, desiccation and starvation tolerance in *D. melanogaster* to contrast variation within and among populations sampled at latitudinal extremes. They found the greatest source of variation in tolerance traits was among genotypes within populations, and only cold resistance showed variation among populations within latitude. Their results, combined with ours, suggest that understanding adaptive potential of metapopulations to global change requires dissecting how adaptive genetic variation is distributed among versus within populations across heterogeneous landscapes. It is therefore necessary to sample a variety of environments within several latitudes at distances where selection could still overcome gene flow. Given these experiments are labour-intensive, studies should either prioritise estimating among-population variation by sampling numerous (>15) populations, or focus on fewer populations from locations that vary in heterogeneity and dissect among versus within population variation. Finally, our populations were maintained at large sizes for c.12 generations, which means it is possible that lab adaptation and genetic drift could have influenced our estimates of among-population variation. However, a study maintaining similarly large populations in the lab found consistent latitudinal clines after c.20 generations (Lasne *et al*. 2018), suggesting this concern should be minimal in our experiment.

### Conclusions

We show that adaptive differentiation across a relatively small scale is comparable to that found across a latitudinal gradient. However, patterns of adaptation changed with treatment temperature and across geography, suggesting that predicting responses to global change needs to consider how adaptation occurs at different geographic scales. Overall, our results support the idea that local adaptation at smaller geographic scales can generate adaptive genetic differentiation that increases the adaptive capacity of populations facing global change. Decisions on managing landscapes should then aim to conserve local-scale environmental heterogeneity to maximise the potential for local adaptation and increase population resilience.

## Acknowledgements

We are grateful to two anonymous reviewers whose constructive comments helped us to greatly improve this paper. We thank Ary Hoffmann for comments on a previous version. We are grateful to Carmen da Silva, Julian Beaman and Belinda van Heerwaarden for helping to sample the latitudinal populations. We thank Andrew Murch for his insights into climatic variation across Victorian wine regions. We also thank Emily Lombardi, Mia Wansbrough and Sandra Hangartner for their help with some of the assays. GMW is grateful to be supported by Australian Research Council fellowships DE200101019 and FT240100466. CMS was supported by Australian Research Council Discovery Project grants DP200102754 and DP220102958.

