## Supplementary material for "Local adaptation in climate tolerance at a small geographic scale contrasts with broad latitudinal patterns"

**Contents**

**Table S1** Sampling locations

**Table S2** Developmental time of each treatment

**Fig. S1** Wing landmarks

**Fig. S2** Quantifying among-population variance

**Table S3** Summary tables for regressions of tolerance traits with mean temperature

**Table S4** Summary tables for regressions of heat tolerance plasticity

**Fig. S3** Correlations among environmental variables

**Methods S1** Establishing populations

**Table S1:** Population locations sampled from across latitude and within Victoria. Rain (mm) is the total rainfall for the four months prior to sampling. Lines represents the number of isofemale lines collected in the field that were used to establish the mass-bred populations. Note that the SD population was created from isofemale lines collected from two wineries in close proximity.

| **Geog-raphy** | **Code** | **State** | **Location** | **Sampling location** | **Latitude** | **Longitude** | **Lines** |
| --- | --- | --- | --- | --- | --- | --- | --- |
| Latitude | INI | Qld | Innisfail | Sellars Bananas | -17.9414 | 146.0553 | 40 |
|  | MAC | Qld | Mackay | FoodPac Fruit | -21.1363 | 148.5098 | 47 |
|  | BAL | NSW | Ballina | Southern Cross Botanicals | -28.7670 | 153.5361 | 19 |
|  | RY | Vic | Yarra Valley | Raynor's Orchard | -37.7924 | 145.5512 | 44 |
|  | TAS | Tas | Huonville | Willie Smith's Organic Cider | -42.9960 | 147.0718 | 29 |
| Victoria | AT | Vic | Daylesford | Atwood Wines | -37.2724 | 144.2296 | 60 |
|  | BE | Vic | Great Western | Best's Winery | -37.1332 | 142.8416 | 48 |
|  | ML | Vic | Mildura | Qualia Winery | -34.2600 | 142.1993 | 60 |
|  | PZ | Vic | King Valley | Pizzini Wines | -36.7886 | 146.4213 | 59 |
|  | SD | Vic | Bairnsdale | Nicholson River Winery | -37.7961 | 147.7565 | 21 |
|  | SD | Vic | Bairnsdale | Tambo Winery | -37.7848 | 147.8202 | 33 |
|  | SH | Vic | Yarra Valley | Long Shadow Wines | -37.7365 | 145.3033 | 50 |
|  | ST | Vic | Rutherglen | Stanton & Killeen Wines | -36.0566 | 146.4227 | 58 |
|  | TH | Vic | Ngambie | Tahbilk Winery | -36.8316 | 145.0851 | 56 |

**Table S2:** Developmental time and replicates of each treatment

| **Temperature** | **Time to eclosion** | **Block 1** | **Block 2** |
| --- | --- | --- | --- |
| 13°C | 36-38 days | 3 egg-picks  21 vials/population | 3 egg-picks  21 vials/population |
| 25°C | 9-10 days | 2 egg-picks  14 vials/population | 3 egg-picks  21 vials/population |
| 29°C | 7-8 days | 2 egg-picks  14 vials/population | 3 egg-picks  21 vials/population |

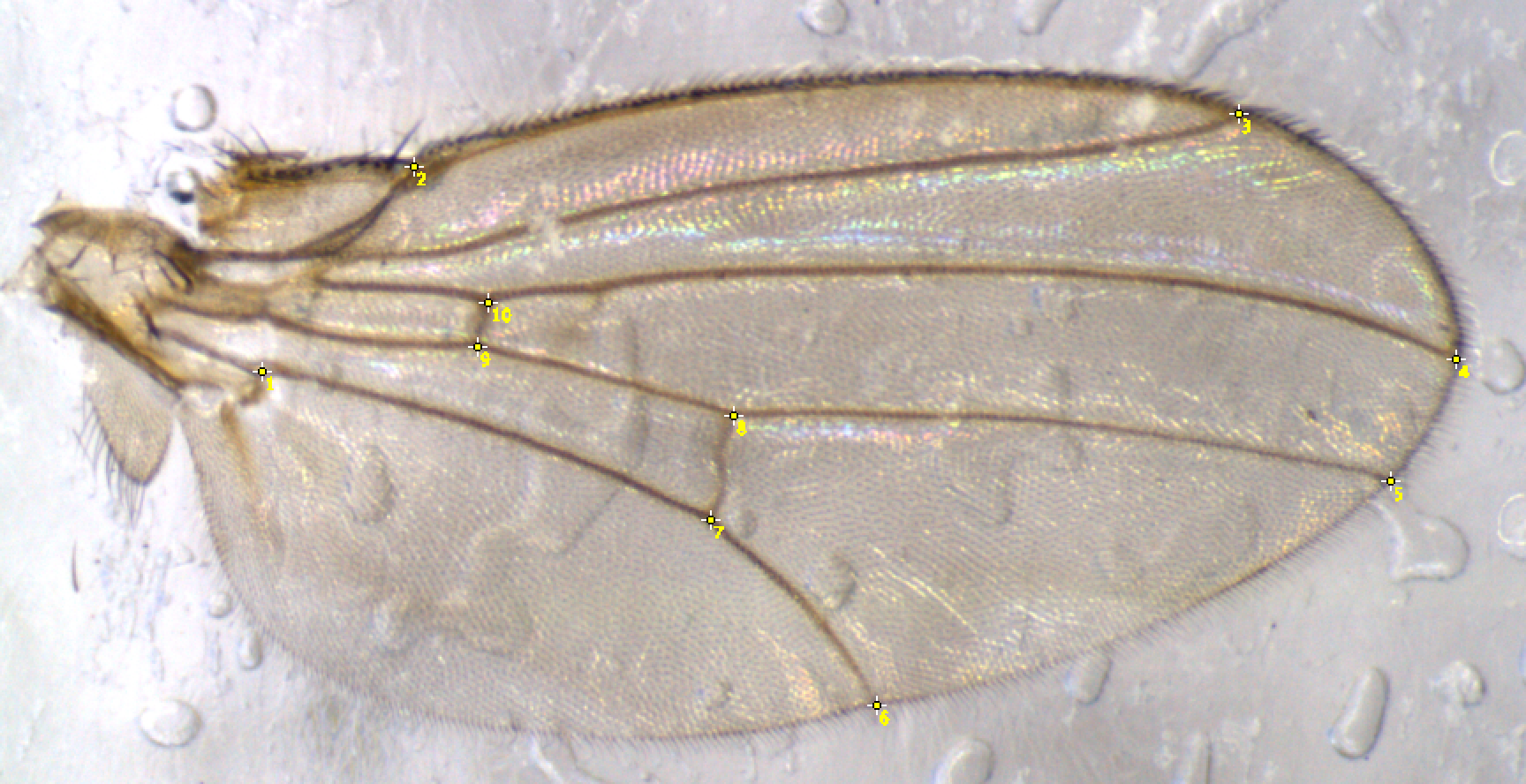

**Fig. S1** Wing landmarks used to estimate centroid size.

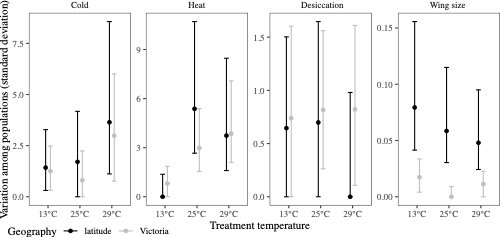

**Fig. S2** Estimates of among-population variance for latitudinal populations (black) versus Victorian populations (grey). Points represent estimates of variance, with credible intervals the 95% confidence interval. These estimates were obtained from equation 1 using the *confint* function.

**Table S3** Summary tables for Type III ANOVA testing the interaction between the environment of origin (O), geography (G; latitude versus Victoria) and treatment temperature (T). Environment includes the mean temperature for the first three traits (cold recovery, heat resistance and wing size), and average daily rainfall for desiccation tolerance. Values in bold are significant at p<0.05.

|  |  | **Cold recovery** | | **Heat resistance** | | **Wing size** | | **Desiccation resistance** | |
| --- | --- | --- | --- | --- | --- | --- | --- | --- | --- |
| **Parameter** | ***df*** | **χ^2^** | **P** | **χ^2^** | **P** | **χ^2^** | **P** | **χ^2^** | **P** |
| Intercept | 1 | 35.89 | **<0.001** | 5.83 | **0.016** | 3033.17 | **<0.001** | 1478.17 | **<0.001** |
| Origin (O) | 1 | 0.35 | 0.554 | 4.52 | 0.033 | 10.32 | **0.001** | 6.84 | **0.009** |
| Geography (G) | 1 | 9.34 | **0.002** | 0.01 | 0.909 | 14.25 | **<0.001** | 18.14 | **<0.001** |
| Treatment (T) | 2 | 20.74 | **<0.001** | 21.27 | **<0.001** | 162.19 | **<0.001** | 8.73 | **0.013** |
| O×G | 1 | 10.53 | **0.001** | 0.00 | 0.994 | 16.76 | **<0.001** | 8.14 | **0.004** |
| O×T | 2 | 1.06 | 0.588 | 18.47 | **<0.001** | 12.11 | **0.002** | 0.70 | 0.705 |
| G×T | 2 | 13.76 | **0.001** | 1.05 | 0.592 | 2.93 | 0.231 | 1.24 | 0.537 |
| O×G×T | 2 | 15.11 | **<0.001** | 1.23 | 0.541 | 2.12 | 0.347 | 0.14 | 0.931 |

**Table S4** Summary Type III ANOVA testing the interaction between temperature **(a)** predictability or **(b)** variation for environment of origin (O), geography (G; latitudinal versus Victorian populations) and treatment temperature (T) on plasticity in heat tolerance. Predictability is quantified as temperature autocorrelation across days for the four months preceding sampling, while variation in temperature is the standard deviation across the entire four months. Values in bold are significant at p<0.05.

|  |  | **(a)** Predictability | | **(b)** Variation | |
| --- | --- | --- | --- | --- | --- |
| **Parameter** | ***df*** | **χ^2^** | **P** | **χ^2^** | **P** |
| Intercept | 1 | 2.30 | 0.129 | 0.39 | 0.534 |
| Autocorrelation (O) | 1 | 3.20 | 0.074 | 1.19 | 0.276 |
| Geography (G) | 1 | **14.77** | **<0.001** | 0.07 | 0.791 |
| Treatment temperature (T) | 1 | 2.64 | 0.104 | **6.79** | **0.009** |
| O×G | 1 | **15.29** | **<0.001** | 0.02 | 0.882 |
| O×T | 1 | 0.11 | 0.736 | 0.03 | 0.864 |
| G×T | 1 | 2.15 | 0.143 | 0.01 | 0.923 |
| O×G×T | 1 | 2.13 | 0.145 | 0.03 | 0.868 |

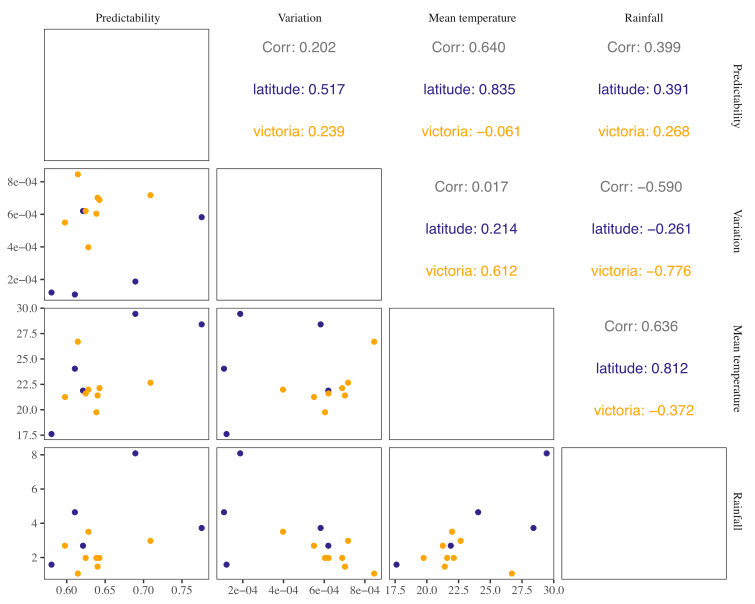

**Fig. S3** Correlations between environmental variables. Predictability is quantified as autocorrelation of mean daily temperature across four months, and variation is the standard deviation of temperature across the four months. Latitudinal populations are represented in blue, Victorian populations in orange and the overall correlation in grey (only above the diagonal). In general, correlations are stronger for latitudinal populations.
**Methods S1** Establishing populations in the laboratory

In the lab, we generated isofemale lines by allowing inseminated females to individually lay for 4 days in 40mL vials containing 7mL of standard yeast-potato-dextrose medium (Brewer’s yeast 36.4g/L; potato flakes 18.2g/L; dextrose 27.3g/L; agar 6.4g/L, nipagin [10% w/v in ethanol] 10.9mL/L; and propionic acid 4.6mL/L), and leaving the larvae to develop at 25°C on a 12:12 hour light:dark cycle. In the following generation, we checked species identification to prevent contamination from the morphologically similar *D. simulans.* In the third generation, we treated each isofemale line with 0.3mg/mL of tetracycline in the food to remove differences among populations in their endosymbionts, particularly *Wolbachia*, which is more prevalent in tropical populations. After three generations, for each population we combined five males with five virgin females from each isofemale line (19–60 lines per population, mean=44.6; **Table S1**) into a mass bred population that we maintained in five 300mL bottles containing 62.5mL of standard food. Each generation, we collected adults and kept them in bottles until they reached peak fertility (c.4 days), after which they were allowed to lay 250–300 eggs in each of the five bottles. After laying, we removed the adults, added cards for pupation and left the larvae to develop.
